# *APOE4* drives widespread changes to the hepatic proteome and alters metabolic function

**DOI:** 10.1101/2025.06.15.659815

**Authors:** Colton R. Lysaker, Chelsea N. Johnson, Vivien Csikos, Edziu Franczak, Maggie Benson, John P. Thyfault, Paige C. Geiger, Jill K. Morris, Heather M. Wilkins

## Abstract

Apolipoprotein E (APOE) is essential for lipid homeostasis and has been extensively studied in the central nervous system, particularly in the context of Alzheimer’s disease (AD). Individuals carrying an *APOE4* allele have an increased risk of AD and exhibit deficits in energy metabolism, including glucose utilization and mitochondrial dysfunction. While the role of APOE in the liver is well characterized, the impact of *APOE* genetic variation on hepatic health and metabolism remains poorly understood. We sought to investigate this using young female and male *APOE3* and *APOE4* targeted replacement mice. We also used *APOE* isogenic induced pluripotent stem cell (iPSC)-derived hepatocyte-like cells (iHLCs) to examine specific effects in a human-relevant cell model. Proteomic and functional assays show that *APOE4* causes extensive changes to liver mitochondrial function in a sex-specific manner in mice and alters glucose and lipid metabolism. *APOE4* also impairs mitochondrial function in iHLCs and shifts metabolism towards glycolysis while modifying expression of extracellular matrix proteins. Additionally, *APOE4* iHLCs display a greater reliance on fatty acids as an energy source and show increased lipid accumulation. Taken together, our findings show that *APOE* genetic variation causes mitochondrial dysfunction and rewires hepatic metabolism.

## Introduction

Apolipoprotein E (APOE) is essential for lipid transport and metabolism. Genetic variation in *APOE* not only influences lipid homeostasis, but also modulates the risk of developing late onset Alzheimer’s disease (LOAD) (1). *APOE* is polymorphic, and variation in *APOE* genotype includes alleles that either confer decreased (*APOE\**ε*2*), neutral (*APOE\**ε*3*), or increased (*APOE\**ε*4*) risk of AD (1, 2). These alleles give rise to the APOE2, APOE3, and APOE4 protein isoforms, which differ in their receptor and lipid-binding preferences (3, 4). APOE2 and APOE3 primarily associate with high-density lipoproteins (HDL), commonly referred to as ‘good cholesterol,’ while APOE4 preferentially binds to low-density lipoproteins (LDL) and very low-density lipoproteins (VLDL), known as ‘bad cholesterols’ (3). Within the human population, *APOE3* is the most common allele with *APOE2* and *APOE4* being less common (5). While there has been considerable focus on how *APOE* genetic variation influences brain health, much remains unknown regarding the liver.

Across the body there are two distinct pools of APOE production. This includes the central nervous system (CNS) and the liver which are separated by the blood brain barrier (BBB) (6, 7). In the periphery, APOE is primarily produced and secreted by hepatocytes (8, 9). Once secreted, APOE binds nascent lipoprotein particles and transports lipids to distal tissues for use (10, 11). The importance of APOE in the liver and other tissues has been highlighted by studies in mice where knockout (KO) of *Apoe* results in significant metabolic stress, specifically dyslipidemia, atherosclerosis, oxidative stress, and inflammation (12–14). One particular study examined mitochondrial proteomic changes in *Apoe* deficient mice and found changes to proteins associated with lipid metabolism and antioxidant response (15). This not only highlights the importance of APOE in broad metabolic function but also its potential effects on mitochondria.

Few studies have examined the role of APOE isoforms on hepatic metabolic health. This includes work with both *in vivo* and *in vitro* models. Of these limited studies, one investigated how APOE isoforms influence hepatic metabolic health in female *APOE* targeted replacement (TR) mice and transformed APOE3 or APOE4 Huh-7 hepatoma cells (16). Their results showed that APOE4 altered liver proteasome activity and decreased adenosine triphosphate (ATP) levels under stress in Huh-7 cells. Despite this, they observed little change to proteins associated with mitochondrial function and autophagy. However, proteomic work in male *APOE* TR mice on a western diet found that APOE4 protects against diet-induced changes to the liver when compared to APOE3 (17). They also showed that *APOE* genotype altered proteins associated with immune response, ER stress, and metabolic function in the liver. Another group found that differences in *APOE* genotype can alter lipidomic profiles in primary human hepatocytes (18). More recently, a serum proteomic analysis in individuals with AD highlighted peripheral protein changes influenced by *APOE* genotype, with several of these proteins secreted by the liver (19).

In this study we investigated how *APOE* genetic variation impacts hepatic metabolic health. We applied proteomic analysis and respirometry to interrogate the effects of *APOE* genotype on hepatic metabolic health using 4-month-old *APOE* TR mice and induced pluripotent stem cell (iPSC)-derived hepatocyte like cells (iHLCs). To our knowledge, this is the first study to leverage iHLCs to investigate the role of *APOE* genotype in hepatic metabolic health *in vitro*. Based on past research, we hypothesized that APOE would drive changes to hepatic metabolism in a genotype dependent manner. Our findings show that *APOE* genetic variation alters the hepatic proteome and modifies metabolic function in both *APOE* TR mice and isogenic iHLCs.

## Methods

### Animals

Male and female homozygous *APOE3* (B6.129P2-*Apoe^tm2(APOE*3)Mae^* N8) and *APOE4* (B6.129P2-*Apoe^tm3(APOE*4)Mae^*N8) targeted replacement (TR) mice were purchased from Taconic Biosciences (Germantown, NY) for this study (n=8 per group for genotype and sex). These *APOE* TR mice were previously generated by removing and replacing the endogenous mouse *Apoe* gene with the human *APOE* gene (20, 21). Mice were maintained on a standard chow diet (Teklad Global Rodent Diet^®^, 8604) for the entire study and housed at ∼25°C with a standard 12-hr light/dark cycle. Mice were aged to 4 months before the administration of 90 mg/kg ketamine and 10 mg/kg xylazine followed by euthanasia. Liver tissue was collected immediately following euthanasia and used for mitochondrial isolation or stored at −80°C. The mice used in this study were also used in another publication (22). Animal work and protocols were conducted under approval from the Institutional Animal Care and Use Committee at the University of Kansas Medical Center.

### Human induced pluripotent stem cell culture

CRIPSR-Cas9 gene edited isogenic human induced pluripotent stem cells (iPSCs) homozygous for *APOE3* or *APOE4* were obtained from The Jackson Laboratory (JAX). These iPSCs were generated as previously described from the KOLF2.1J parental male cell line (23). Two pairs of isogenic iPSCs were used in this study, with Pair A consisting of JIPSC001162 (*APOE3* REV/REV) and JIPSC001150 (*APOE4* SNV/SNV), and Pair B consisting of JIPSC001268 (*APOE3* REV/REV) and JIPSC001142 (*APOE4* SNV/SNV). Cells were cultured with mTeSR™ Plus (STEMCELL Technologies) and maintained on CellAdhere™ Laminin 521 (LN521) (STEMCELL Technologies) matrix coated culture plates (TPP). iPSCs were passaged using ReLeSR™ (STEMCELL Technologies), once they reached approximately 70-90% confluency. All iPSCs used in experiments were kept below passage 20.

### Generation of hepatocyte-like cells

Differentiation of iPSCs to hepatocyte-like cells (iHLCs) was carried out using two previously described protocols with minor modifications (24, 25). Basal medium (BDM) for differentiation was generated by combining RPMI 1640 (Gibco) with B-27™ supplement (Gibco) and penicillin-streptomycin (final concentration 5 U/mL and 5 μg/mL). This was then used for subsequent differentiation medias.

On day 1, nearly confluent iPSCs were passaged onto LN521 coated culture plates (TPP) at a density of 2.5×10^4^ cells/cm^2^ in mTeSR™ Plus with 10 μM Y-27632 ROCK inhibitor (STEMCELL Technologies). The following day media was substituted with fresh definitive endoderm (DE) media which contained BDM supplemented with 100 ng/mL Activin A (MedChemExpress) and 3 µM CHIR99021 (STEMCELL Technologies). Fresh DE media was replaced daily until day 5, after which the cells were cultured in hepatic endoderm (HE) media consisting of BDM supplemented with 5 ng/mL basic fibroblast growth factor (bFGF) (MedChemExpress), 20 ng/mL bone morphogenetic protein 4 (BMP4) (MedChemExpress), and 0.5% DMSO. Full media changes were performed daily with HE media until day 10. Cells were then cultured for 5 days with full media changes in immature hepatocyte (IMH) media containing BDM plus 20 ng/mL hepatocyte growth factor (HGF) (MedChemExpress) and 0.5% DMSO. Finally, on day 15, the media was replaced with mature hepatocyte (MH) medium, followed by full media changes daily for another 10 days. MH media consisted of hepatocyte basal media (Lonza), SingleQuots® supplements (Lonza), 20 ng/mL hepatocyte growth factor (HGF) (MedChemExpress), 20 ng/mL Oncostatin M (MedChemExpress), 100 nM dexamethasone (Sigma), and 0.5% DMSO. iHLCs used in experiments were between days 20-25 of differentiation.

For downstream functional assays, iHLCs were replated on day 20 of differentiation. Mature iHLCs were washed with 1X PBS (Gibco) and then dissociated with pre-warmed (37°C) TrypLE (Gibco) for 10-15 min. Cells in TrypLE were then diluted with DMEM/F-12 and passed through a 70 μm strainer. Cells were centrifuged for 5 min at 300 x g, resuspended in MH media containing 10 μM Y-27632 ROCK inhibitor (STEMCELL Technologies), and plated on LN521 coated plates. Plating density varied depending on downstream assays.

### Mitochondrial isolation from whole liver

Mitochondria were isolated from fresh mouse liver samples as previously described (26, 27). Briefly, livers from mice were immediately placed in 8 mL of ice-cold mitochondrial isolation buffer (MIB) (220 mM mannitol, 70 mM Sucrose, 10 mM Tris, 1 mM EDTA, pH 7.4) and homogenized using a Teflon pestle on ice. Homogenates were then centrifuged at 1,500 x g for 10 min at 4°C to remove cell debris and fat. To remove additional debris, the supernatant was collected and passed through a gauze filter before being centrifuged at 8000 x g for 10 min at 4°C. The pellet was then resuspended on ice in 6 mL of MIB with a glass dounce homogenizer before being centrifuged again at 6,000 x g for 10 min at 4°C. Resuspension of the new pellet was performed on ice in 4 mL of MIB plus 0.1% BSA using a glass dounce homogenizer, followed by centrifugation at 4,000 x g for 10 min at 4°C. The final pellet containing mitochondria was resuspended in 500 μL of ice-cold MiR05 buffer (0.5 mM EGTA, 3mM MgCl_2_, 60 mM KMES, 20 mM glucose, 10 mM KH_2_PO_4_, 20 mM HEPES, 110 mM sucrose, 0.1% BSA, pH 7.1). A BCA assay (ThermoFisher) was performed to determine protein content. Mitochondrial samples were then either immediately used for respiration analysis or frozen at −80°C.

### Proteomics

Preparation of proteomic samples was completed as described below. Whole liver samples were lysed with RIPA buffer (Millipore) plus protease inhibitors (ThermoFisher, A32965). Tissue lysates were then centrifuged at maximum speed for 10 min before the supernatant was transferred to a new tube. A BCA assay was performed to quantify protein concentration. Samples were then transferred to a new tube and diluted with fresh RIPA to 1 μg/μL (100 μg total). Protein content was determined from previously frozen isolated liver mitochondria samples which were diluted to 1 μg/μL (100 μg total).

iHLCs were harvested for proteomics on day 21 of differentiation. Proteomic analysis was performed on two independent differentiation experiments using Pair A isogenics, referred to as Batch 1 and Batch 2. Plates containing iHLCs were placed on ice before media was removed and discarded. Cells were then washed once with ice cold 1X PBS and then scraped in 1X PBS containing protease inhibitors (ThermoFisher, A32965). Cells were collected into 1.7 mL tubes and centrifuged at maximum speed for 5 min. The supernatant was removed, and the pellet was resuspended and lysed in 500 uL RIPA (Millipore) with protease inhibitors (ThermoFisher, A32965). A BCA assay was used to determine protein content and samples were then diluted to 1 μg/μL (100 μg total) in fresh RIPA buffer.

Samples were then sent to the University of Arkansas Medical Center (UAMS) for proteomics. Briefly, protein samples were extracted using chloroform/methanol phase separation with a subsequent trypsin digestion to acquire peptides. Samples were then analyzed using liquid chromatography mass spectrometry (LC/MS) on a Orbitrap Exploris™ 480 (ThermoFisher) with data independent acquisition (DIA).

### APOE enzyme-linked immunosorbent assay

Quantification of APOE from mouse livers was completed using a commercially available ELISA kit (ThermoFisher, EHAPOE). Mouse livers were dounce homogenized on ice in RIPA buffer (Millipore) with the addition of protease inhibitors (ThermoFisher, A32965). Liver homogenates were then centrifuged at 12,500 x g for 10 min. Supernatants were collected in new tubes and diluted 1:100 in assay diluent buffer. The ELISA was completed per the manufacturer’s instructions. Final concentrations were normalized to protein determined using a BCA assay (ThermoFisher).

### Western blotting

Lysates from cells were collected on ice and lysed using RIPA buffer (Millipore) plus protease inhibitors (ThermoFisher, A32965). A BCA assay (ThermoFisher) was performed, and an equal amount of protein was resolved on 4-15% Criterion TGX gels (Bio-Rad). Gels were transferred to PVDF membranes (Bio-Rad) and blocked overnight at 4°C with 5% bovine serum albumin (BSA) in 1X phosphate buffered saline with Tween (PBST). The albumin western blot was blocked with animal free blocking solution (Vector Laboratories, SP-5030-250). Primary antibodies diluted at 1:1000 were applied to the membranes and incubated overnight at 4°C. Blots were washed three times in 1X PBST and incubated at room temperature for 1 hr with secondary antibodies diluted 1:2000 in 5% non-fat dry milk (NFDM) prepared in 1X PBST. Blots were then imaged using SuperSignal West Femto or West Dura ECL substrates (ThermoFisher) on a ChemiDoc XRS imaging system (Bio-Rad). Amido black (Sigma) was used to stain total protein. Primary antibodies included ALB (Abcam, ab207327), APOE (Abcam, ab183597), ASGR1 (Abcam, ab254261), HNF4A (Abcam, ab92378), NANOG (Abcam, ab109250), OCT4 (Abcam, ab181557), and SOX17 (Abcam, ab224637).

### Immunocytochemistry

iHLCs on day 23 of differentiation were fixed using 4% paraformaldehyde (PFA) for 15 min. Cells were washed three times with 1X PBS (Gibco) before being permeabilized with 0.1% Triton X-100 (Sigma) for 10 min in 1X PBST. Cells were washed three times with 1X PBS (Gibco) and blocked for 1 hr at room temperature with 1% BSA in 1X PBST. The blocking solution was removed, and cells were incubated overnight with primary antibodies at 4°C. The following day cells were washed with 1X PBS (Gibco) three times followed by a 1 hr incubation at room temperature with secondary fluorophore conjugated antibodies. The nuclear counterstain Hoechst was used at a concentration of 2 μg/mL for 30 min. Finally, cells were washed three times with 1X PBS (Gibco) before being imaged on a BioTek Cytation 1 (Agilent). Primary antibodies included APOE diluted 1:400 (Abcam, ab183597), ASGR1 diluted 1:200 (Abcam, ab254261), and HNF4A diluted 1:200 (Abcam, ab92378). We used DyLight™ 488 conjugated secondary antibody (ThermoFisher) at a concentration of 2 μg/mL.

### Isolated mitochondria respiration

Liver mitochondrial oxygen consumption rates (OCR) were measured using a Seahorse XFe96 Bioanalyzer (Agilent). Isolated mitochondria were diluted to 15-20 µg per 180 µL using mitochondrial assay solution (MAS) (220 mM Mannitol, 70 mM Sucrose, 10 mM K_2_HPO_4_, 5 mM MgCl_2_, 2 mM HEPES, 1 mM EGTA, 0.2% fatty acid-free BSA). Mitochondria were loaded into Seahorse cell culture plates at 180 µL per well. The Seahorse plate was centrifuged at 1000 x g for 5 min with the brake set at 5. State 2 respiration rates were determined using palmitoyl-coenzyme A (PCoA, 10 µM) or 5 mM pyruvate and 5 mM malate. State 3 respiration was acquired after the addition of 4 mM ADP. Succinate was then injected at a concentration of 10 mM to acquire state 3S. The final addition of carbonyl cyanide-p-trifluoromethoxy phenylhydrazone (FCCP, uncoupled, 4 µM) assessed maximal respiratory flux. Respiration values were normalized to total protein amount loaded and across days of mitochondrial respiration analysis.

### Cellular respiration

A Seahorse XF Pro Analyzer (Agilent) was used to determine iHLC OCR and extracellular acidification rates (ECAR). Cells were plated at a density of 35,000 per well on LN521 coated Seahorse cell culture plates. Substrates were successively injected to measure OCR or ECAR for different respiration rates*. Mitochondrial Stress Test*: Seahorse base medium contained 10 mM glucose, 2 mM glutamine, with injections A) 2 µM oligomycin B) and C) 0.5 µM FCCP D) 1 µM antimycin A and rotenone. *Glycolysis Stress Test:* Seahorse base medium contained 2 mM glutamine with injections A) 10 mM glucose B) 2 µM oligomycin C) 100 mM 2-deoxy-glucose (2-DG). *Electron Transport Chain Flux:* Medium was 1X MAS supplemented with 10 mM pyruvate/5 mM malate/4 mM ADP and 1 nM plasma membrane permeabilization (PMP) reagent with injections A) 10 mM Succinate/ 1 µM rotenone B) 1 µM antimycin A C) 0.5 mM TMPD/ 1 mM ascorbate D) 50 mM azide. *Glucose/Pyruvate and LCFA Substrate Oxidation Stress Tests*: To determine substrate oxidation we used Seahorse base medium containing 10 mM glucose, 2 mM glutamine, and 1 mM pyruvate with the following injection scheme: A) assay medium or 2 µM UK5099 or 4 µM Etomoxir B) 2 µM oligomycin C) 0.5 µM FCCP D) 1 µM antimycin A and rotenone.

OCR and ECAR were measured following a 3-min mix after the addition of injections and measured over 3 min three times. All data were analyzed using the Agilent Seahorse Wave software and Microsoft Excel.

### LipidTOX imaging and quantification

To quantify lipid droplets (LDs) in iHLCs, we used HCS LipidTOX™ Red neutral lipid stain (H34476, Thermo Fisher Scientific). LD staining of iHLCs was done as specified in the manufacturer’s instructions. Cells were replated at a density of 3.0×10^4^ cells/well of a transparent bottom 96-well plate (204626-100, Agilent) and allowed to recover for one day. Cells were fixed with 4% PFA for 20 min at room temperature and then washed three times with 1X PBS (Gibco). 1X LipidTOX stain was subsequently added with Hoechst at a concentration of 2 μg/mL, and cells were incubated for 30 min at room temperature before imaging on a BioTek Cytation 1 (Agilent). Data was acquired and analyzed using BioTek Gen5 software (Agilent). We quantified the average number of LDs per cell in each field of view at 20X magnification using a spot counting feature. We also quantified the average size (diameter) of LDs. Parameters for analysis included a cell size diameter cutoff of 30 μm and LD size cutoff of 0.2-5 μm.

### Mitochondrial indicator assays

iHLCs were plated at a density of 2.0-3.0×10^4^ cells/well of a transparent bottom 96 well plate (3603, Corning) and allowed to recover for one day. To assess mitochondrial membrane potential we used 200 nM tetramethyl rhodamine, ethyl ester (TMRE) (ThermoFisher). Mitochondrial superoxide was measured with 500 nM MitoSOX™ Red (ThermoFisher) and hydrogen peroxide (H_2_O_2_) was determined using 50 μM Amplex™ Red (ThermoFisher) with 0.1 U/mL horseradish peroxidase (HRP). Levels of mitochondrial calcium (Ca^2+^) were measured using 1 μM Rhod-2, AM (ThermoFisher). Hoechst (ThermoFisher) was used for normalization at a concentration of 10 μg/mL. Cells were stained with respective dyes diluted for 30 min at 37°C and then washed twice with 1X Hanks’ Balanced Salt Solution (HBSS) (Gibco). Fluorescence intensity for each dye was read with a Tecan Infinite 200 PRO^®^ and normalized to Hoechst. Representative images for TMRE were acquired at 20X magnification using a BioTek Cytation 1 (Agilent).

### Proteomic data analysis

Data analysis of samples was completed using Spectronaut® software (Biognosys version 18.3). UniProt databases were used to identify both mouse (*Mus musculus*) and human (*Homo sapien*s) proteins. Quality control (QC) and normalization was carried out with proteiNorm (28). MitoCarta 3.0 was used to further identify mitochondrial specific proteins in the isolated liver mitochondria samples (29). Proteins with a p-value < 0.05 were considered significant. QIAGEN Ingenuity Pathway Analysis (IPA) was used for pathway, upstream regulator, and comparison analysis. Proteins included in pathway analysis had a p-value cutoff of < 0.2 for whole liver and isolated mitochondria samples, and < 0.05 for iHLCs. Pathways with a z-score of ≤ −2 and ≥ +2 and a log fold change (logFC) of ≥ 1.3 were considered significant. Proteins involved in specific pathways or cellular functions were identified by cross referencing protein lists with multiple databases including Kyoto Encyclopedia of Genes and Genomes (KEGG), Search Tool for the Retrieval of Interacting Genes/Proteins (STRING), and IPA. STRING analysis was also used to show selected protein interactions.

### Statistical analysis

Values are shown in figures as mean ± SD. Data analysis was completed using GraphPad Prism 10 software and Microsoft Excel. Outliers were identified and removed from datasets using Grubbs’ test with a significant alpha of 0.05. Significance for simple comparisons was calculated using an unpaired t test. For comparisons of multiple groups, we used a two-way analysis of variance (ANOVA) followed by Fisher’s Least Significant Difference (LSD) post-hoc test if significant interactions were identified. A cutoff p-value < 0.05 was considered significant for all analyses. Pearson correlation analysis was used to assess correlations. All graphs and figures were generated using GraphPad Prism 10 software and BioRender.com.

## Results

### APOE genotype and sex alter the hepatic proteome in young mice

Four-month-old male and female *APOE3* or *APOE4* targeted replacement (TR) mice (n=8 per genotype/sex) were used to examine whole liver and isolated mitochondrial proteomics in addition to seahorse mitochondrial function assays (**Figure 1A**). We first examined whole liver protein expression of APOE between male and female *APOE3* and *APOE4* mice and observed no difference at 4 months (**Figure 1B**). Whole liver proteomics revealed 402 differently expressed (DE) proteins between female *APOE4* and *APOE3* mice versus 58 between male *APOE4* and *APOE3* mice (**Figure 1C**). Female *APOE4* TR mice when compared to female *APOE3* TR mice had increased expression of proteins involved in mitochondrial function and bioenergetics (Ndufaf3, Fdft1, Scd1, Ado, Gnpnat1) and peroxisomal function (Pex5) with reduced expression of proteins involved in cholesterol metabolism (Acat2), glycolysis/gluconeogenesis (Aldob), mitochondrial ribosomal function (Mrps34, Mrpl53), mitochondrial/ER interactions/autophagy (Sigmar1), and cell signaling (Plcb3) (**Figure 1D**). IPA revealed significant upregulation of mitochondrial protein degradation, electron transport in the respiratory chain/oxidative phosphorylation, mitochondrial protein import, and mechanistic target of rapamycin (mTOR) signaling pathways in *APOE4* female TR mice when compared to their *APOE3* counterparts (**Figure 1E**). Down regulated pathways included protein ubiquitination, glucose metabolism, mitotic metaphase and anaphase, ribosomal quality control, and roundabout (ROBO) signaling pathways in *APOE4* female mice (**Figure 1E**).

**Figure 1.**
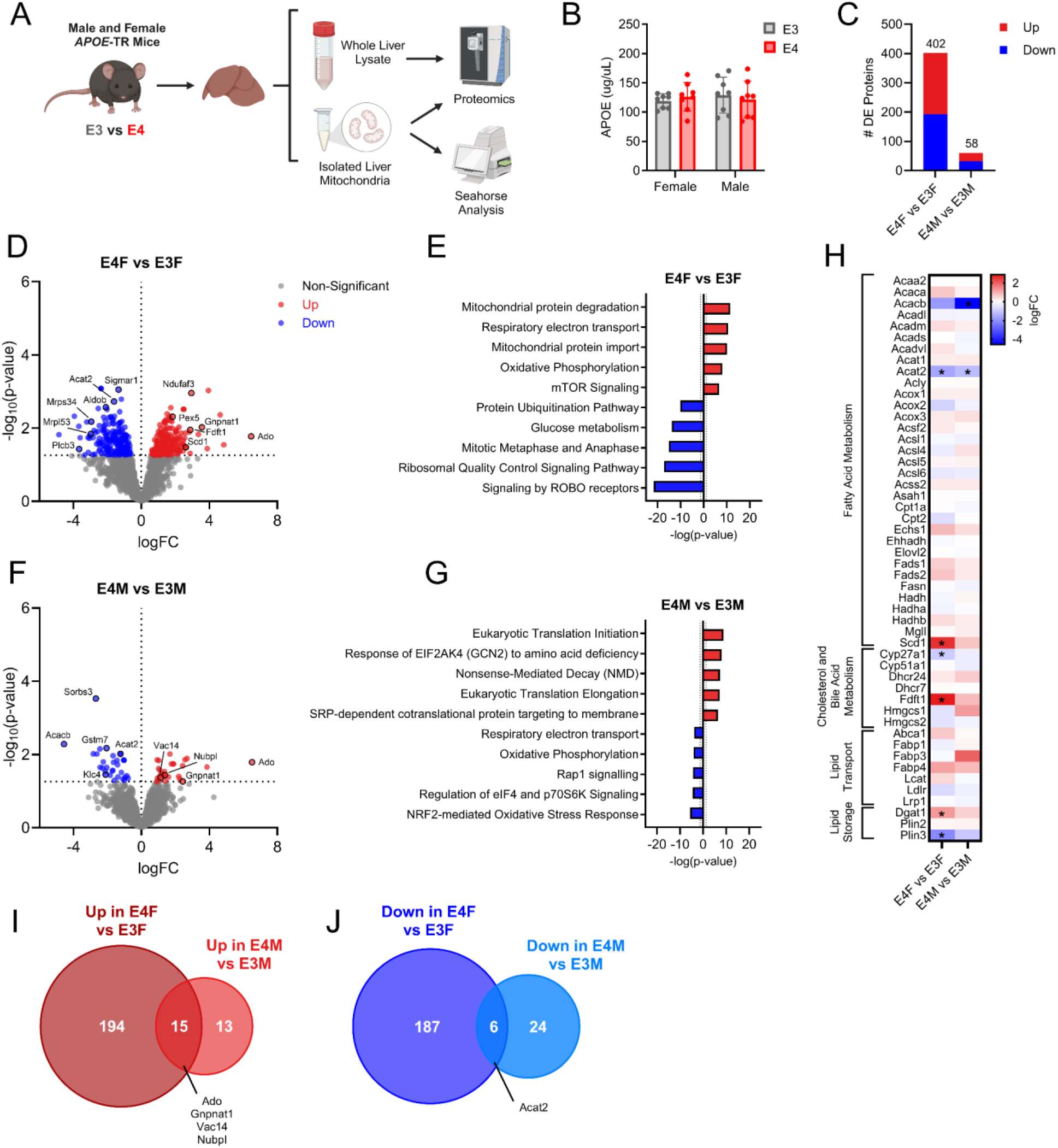
Whole liver proteomics in male and female *APOE3* and *APOE4* mice. A. Study schematic showing primary outcomes in *APOE* targeted replacement (TR) mice. B. APOE protein expression from whole liver measured via ELISA in male and female *APOE3* and *APOE4* mice, n=8 per group. C. The number of differentially expressed (DE) proteins in male and female *APOE3* and *APOE4* mice. D. Volcano plot showing up and down regulated proteins in female *APOE4* vs. *APOE3* mice. E. IPA pathway showing top 5 up and down regulated pathways in female *APOE4* vs. *APOE3* mice. F. Volcano plot showing up and down regulated proteins in male *APOE4* vs. *APOE3* mice. G. IPA pathway analysis showing top 5 up and down regulated pathways in male *APOE4* vs. *APOE3* mice. H. Heatmap of proteins involved in fatty acid metabolism, cholesterol and bile acid metabolism, lipid transport and lipid storage showing up and down regulated proteins between *APOE4* and *APOE3* mice. I. Venn diagram showing the number of shared upregulated proteins between comparisons of female *APOE4* vs. *APOE3* mice and male *APOE4* vs. *APOE3* mice. J. Venn diagram showing the number of down regulated proteins shared between comparisons of female *APOE4* vs. *APOE3* mice and male *APOE4* vs. *APOE3* mice. n=3 for proteomic outcomes. Data are shown as mean ±SD.

Male *APOE4* mice had increased expression of proteins involved in mitochondrial function and bioenergetics (Ado, Nubpl, Gnpnat1) and lysosomal function (Vac14) with reduced expression of proteins involved in cholesterol metabolism (Acat2, Acacb), antioxidant defense (Gstm7), and autophagy or microtubule/actin signaling (Sorbs3, Klf4) (**Figure 1F**). IPA revealed a significant upregulation of eukaryotic translation initiation, amino acid deficiency, nonsense mediated decay, eukaryotic translation elongation, and signal recognition particle (SRP)-dependent co-translational protein targeting pathways in male *APOE4* mice (**Figure 1G**). Down regulated pathways included oxidative phosphorylation, Rap1 signaling, regulation of eIF4/p70S6K, and NRF2 oxidative stress response pathways in *APOE4* male mice (**Figure 1G**).

Sex-specific differences were observed irrespective of *APOE* genotype in the mice (**Figure S1**). Of note, *APOE3* male mice showed an increase in peroxisomal lipid metabolism when compared to their female counterparts (**Figure S1D**). To examine sex-specific differences in lipid proteins, we generated a heat map of fatty acid metabolism, cholesterol metabolism, and lipid-related pathways. Further analysis of fatty acid proteins between mice revealed decreased expression in both male and female *APOE4* mice for Acat2 when compared to their *APOE3* counterparts, with Acacb having decreased expression in male *APOE4* mice only (**Figure 1H**). Furthermore, female *APOE4* mice had upregulation of Scd1 (**Figure 1H**). Cholesterol and bile acid metabolism revealed decreased Cyp27a1 expression in *APOE4* female mice with an upregulation Fdft1 (**Figure 1H**). Lipid storage pathway protein Dgat1 was upregulated while Plin3 was down regulated in female *APOE4* mice (**Figure 1H**). Female and male mice shared 15 upregulated proteins and 6 down regulated proteins when comparing between *APOE4* and *APOE3* mice (**Figure 1I and J**). These findings highlight the sex-specific effects of *APOE* genotype on liver proteomics.

### Mitochondrial proteins and pathways are modified by APOE genotype and sex

We next examined isolated mitochondrial proteomics from *APOE* TR mice liver samples. Before any analysis was performed, we first cross reference the isolated mitochondria protein library with MitoCarta 3.0 (29). Female *APOE4* mice had increased expression of fatty acid beta oxidation (FAO) proteins (Zadh2 and Echdc1), and proteins involved in regulating reactive oxygen species (ROS), electron transport and oxidative phosphorylation (Cisd3) (**Figure 2A**). The mitochondrial dynamic protein involved in fusion Mfn1, FAD production protein Flad1, and mitochondrial protease Yme1l1 were down regulated in *APOE4* female mice (**Figure 2A**). Male *APOE4* mice had increased expression of DNA repair proteins (Oxnad1), mitochondrial calcium homeostasis proteins (Slc25a23), and proteins involved in regulating ROS, electron transport and oxidative phosphorylation (Cisd3) (**Figure 2B**). Mitochondrial complex I proteins Ndufa3 and Ndufb3, apoptotic regulator Hax1, and complement protein C1qbp were down regulated in male *APOE4* mice (**Figure 2B**). Female *APOE4* mice showed an upregulation in the mitochondrial dysfunction pathway but a down regulation in mitochondrial protein import, complex III assembly, TCA cycle, and glyoxylate metabolism and glycine degradation pathways (**Figure 2C**). Male *APOE4* mice showed an upregulation in granzyme A signaling and TCA cycle pathways, but a down regulation in complex IV assembly, complex I biogenesis, oxidative phosphorylation, and respiratory electron transport pathways (**Figure 2D**). Mitochondrial protein import and protein degradation pathways showed differences between whole liver and isolated mitochondrial proteomics in female mice, where Chchd2, Mtx1, Ssbp1, and Yme1l1 were differentially expressed between whole liver and isolated mitochondria (**Figure 2E**). Sex-specific differences were observed in proteins that are involved in the TCA cycle and pyruvate metabolism, where females showed a down regulation of these proteins compared to males between *APOE4* and *APOE3* mice (**Figure 2F**). Complex I, II, III, IV, and V components also had sex-specific expression patterns between *APOE4* and *APOE3* mice (**Figure 2G**). Sex-specific differences were observed irrespective of *APOE* genotype in these mice (**Figure S2**). This included a decrease in proteins and pathways associated with the TCA cycle in male *APOE3* mice when compared to females (**Figure S2C**).

**Figure 2.**
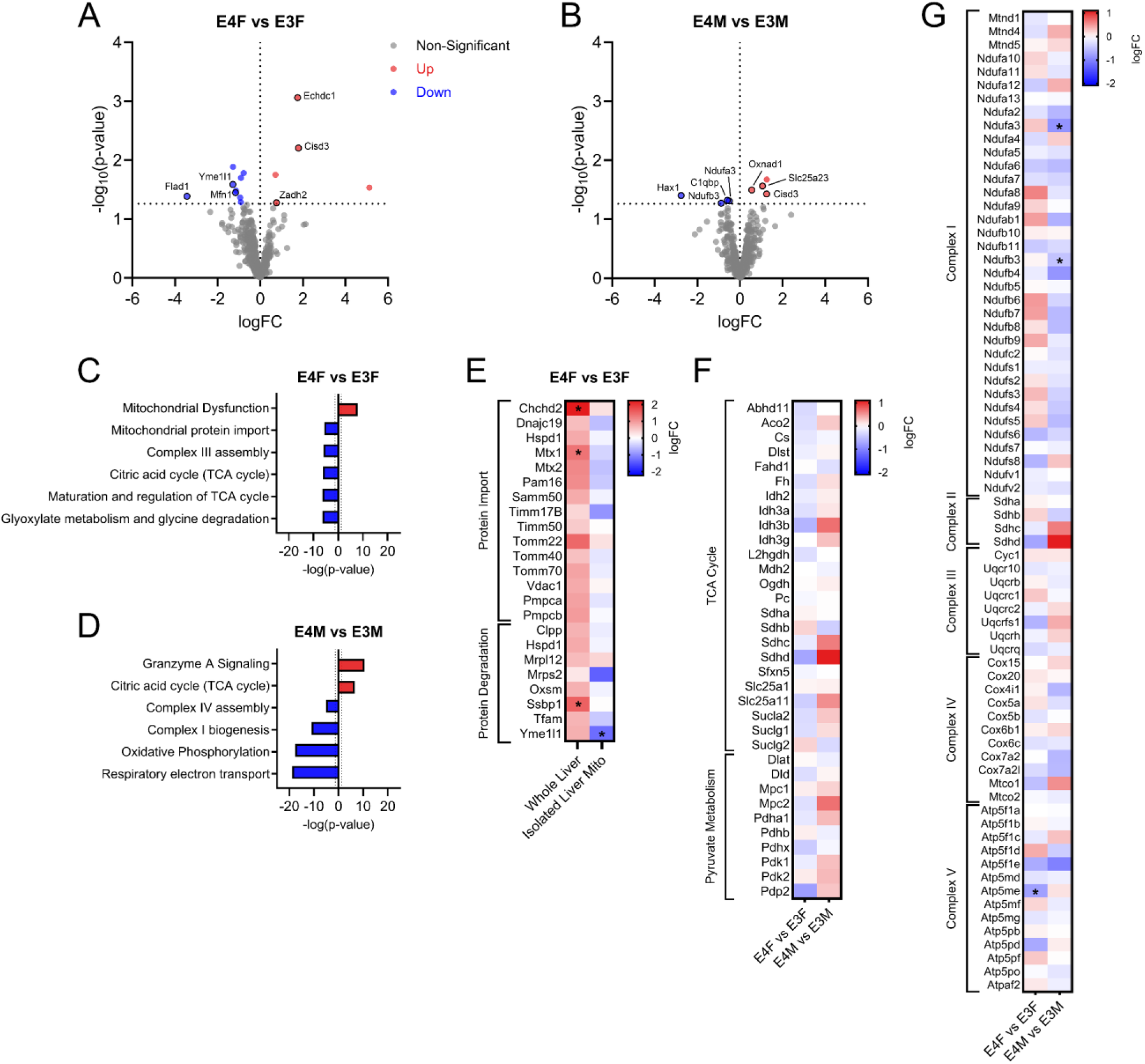
Isolated liver mitochondrial proteomics in male and female *APOE3* or *APOE4* mice. A. Volcano plot showing up and down regulated proteins between female *APOE4* and *APOE3* mice. B. Volcano plot showing up and down regulated proteins between male *APOE4* and *APOE3* mice. C. IPA pathway analysis showing top up and down regulated pathways between female *APOE4* and *APOE3* mice. D. IPA pathway analysis showing top up and down regulated pathways between male *APOE4* and *APOE3* mice. E. Heatmap comparing whole liver to isolated mitochondria results of proteins involved in mitochondrial protein import and degradation between *APOE4* and *APOE3* female mice. F. Heatmap of proteins involved in TCA cycle and pyruvate metabolism between *APOE4* and *APOE3* female and male mice. G. Heatmap of oxidative phosphorylation complex I-V components between *APOE4* and *APOE3* female and male mice. n=3 for proteomic outcomes.

### APOE genotype and sex interact to alter mitochondrial respiration in the liver

To examine mitochondrial function, we leveraged Seahorse respirometry analysis in isolated liver mitochondria from *APOE* TR mice. Fatty acid driven respiration (PCoA) showed a main effect of sex for state 2 and state 3 respiration with increased respiration in males (**Figure 3A and B**). We also observed a trend for increased state 2 and state 3 fatty acid driven respiration in *APOE4* mice when compared *APOE3* mice (**Figure 3A and B**). Fatty acid driven state 3S and uncoupled respiration showed main effects of sex and an interaction between sex and genotype, where male *APOE4* mice had reduced uncoupled respiration (**Figure 3C and D**). Carbohydrate driven respiration showed no differences between sex or genotype for state 2 or state 3 respiration (**Figure 3E and F**). Carbohydrate driven state 3S and uncoupled respiration showed a main effect for an interaction between sex and genotype, where female *APOE4* mice had increased respiration for these states when compared to *APOE3* female mice (**Figure 3G and H**). Additionally, male *APOE3* mice had increased respiration for these states when compared to their female counterparts (**Figure 3G and H**).

**Figure 3.**
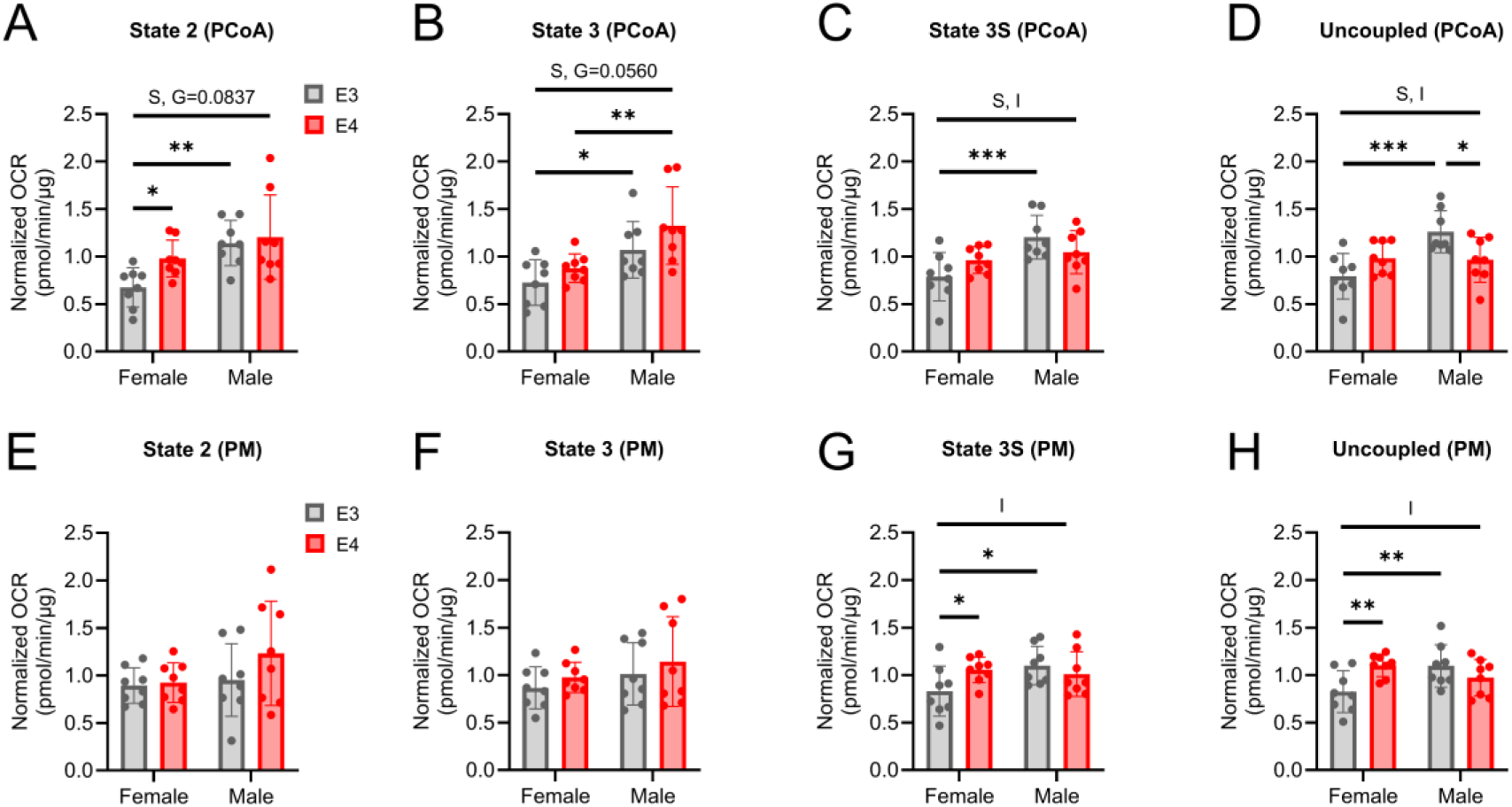
Mitochondrial respiration from isolated liver mitochondria in male and female *APOE3* and *APOE4* mice. Fatty acid (PCoA) driven respiration at A. State 2 B. State 3 C. State 3S and D. Uncoupled respiration states. Carbohydrate (PM) driven respiration at A. State 2 B. State 3 C. State 3S and D. Uncoupled respiration states. n=8 per genotype and sex. Data are shown as mean ±SD. *p<0.05, **p<0.01, ***p<0.001. S=sex, G=genotype, I=interaction.

### APOE genotype drives significant changes to the proteome in iHLCs

To examine potential hepatocyte specific effects of *APOE* genotype, we generated induced pluripotent stem cell (iPSC)-derived hepatocyte-like cells (iHLC) from two isogenic pairs of iPSCs. Characterization of these iHLCs showed expected expression of mature hepatocyte (MH) markers (**Figure S3**). Our primary goal was to examine proteomic and bioenergetic outcomes in these cells (**Figure 4A**). We carried out proteomic analysis on iHLCs generated in two batches from isogenic Pair A, to control for potential variability in differentiation efficiency (**Figure S3**). APOE protein expression in iHLCs was the same for *APOE3* and *APOE4* derived cells in Batch 1 but decreased in *APOE4* iHLCs in Batch 2 (**Figure 4B**). Differences between batches are likely a consequence of higher mature hepatocyte protein expression in Batch 1 when compared to Batch 2, which may affect APOE expression patterns (**Figure S3**) (30). *APOE4* iHLCs from Batch 1 showed 1258 upregulated proteins and 902 down regulated proteins when compared to *APOE3* cells, while Batch 2 *APOE4* iHLCs showed 1097 upregulated proteins and 1115 down regulated proteins (**Figure 4C and D**). Proteins that were upregulated in *APOE4* iHLCs in both batches included those associated with glycolysis, inflammation, and extracellular matrix (ECM) remodeling, while those down regulated in both batches were associated with ribosomal function, cell proliferation, and regeneration (**Figure 4C-E**). We also observed differential expression of proteins associated with AD. This included decreased amyloid precoursor protein (APP) levels in *APOE4* iHLCs (**Figure 4E**). IPA analysis of pathways showed an upregulation in neddylation, Golgi/ER trafficking, autophagy, cGAS-STING, and glycolysis pathways among others in *APOE4* iHLCs (**Figure 4F**). Down regulated pathways in both batches included mitochondrial translation, oxidative phosphorylation, spliceosome, processing of capped intron less pre-mRNA, nucleotide excision repair, complex I biogenesis, and transcription in *APOE4* iHLCs (**Figure 4F**). Due to differences associated with ECM remodeling, we next performed STRING and Gene Ontology (GO) analysis of top differentially expressed proteins from **Figure 4E**. Interestingly, cellular components associated with the ECM were significantly enriched in *APOE4* iHLCs (**Figure 4G, H**).

**Figure 4.**
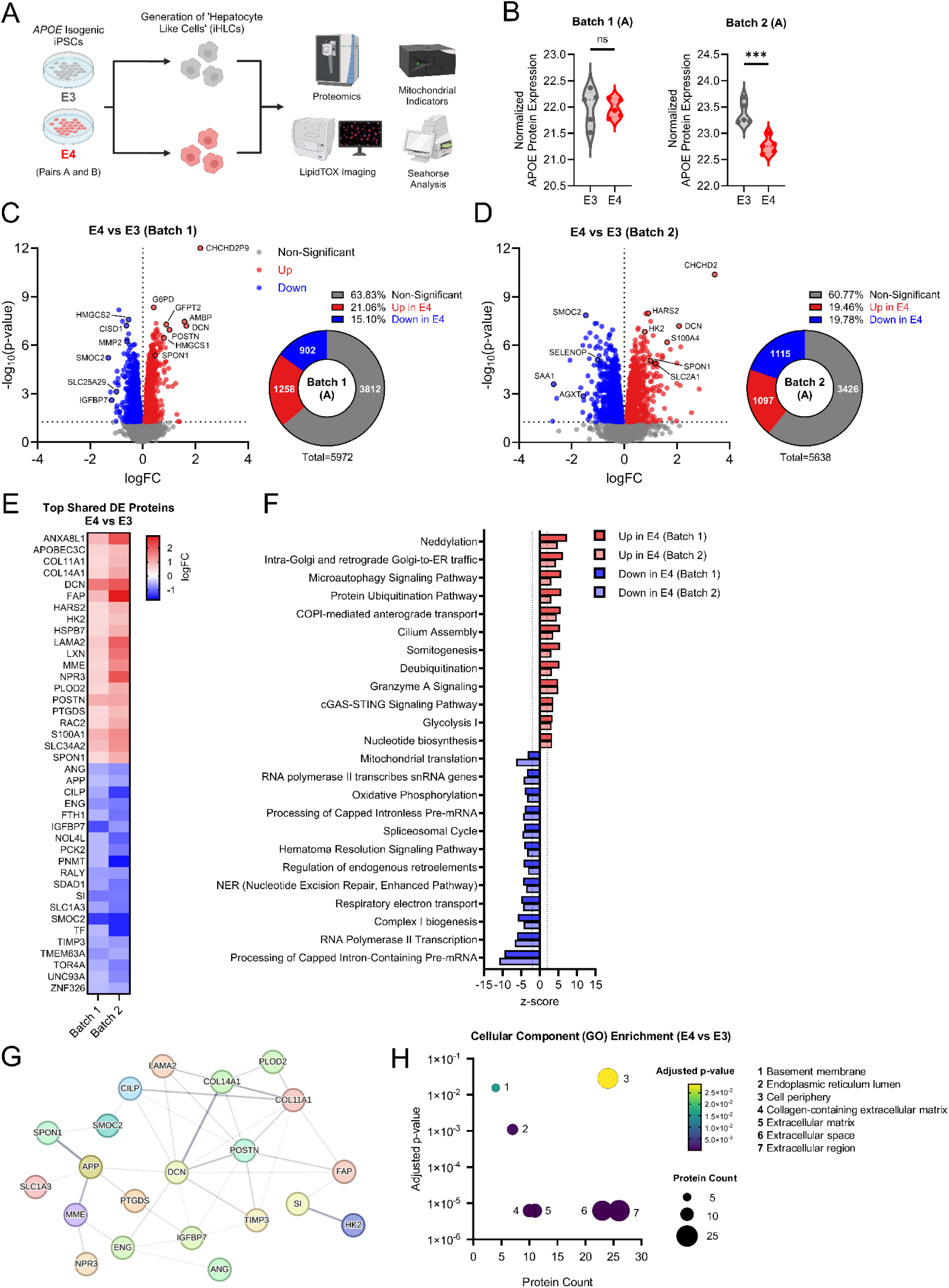
Proteomics analysis of *APOE* isogenic iHLCs. A. iHLC study schematic. Two pairs of isogenic iPSCs (A and B) homozygous for either *APOE3* or *APOE4* were used to generate iHLCs and examine proteomic and bioenergetic outcomes. B. APOE protein expression between two sample batches in Pair A isogenics. C. Volcano plot showing up and down regulated proteins between *APOE4* and *APOE3* iHLCs from Batch 1. D. Volcano plot showing up and down regulated proteins between *APOE4* and *APOE3* iHLCs from Batch 2. E. Heatmap of the top 20 most significant upregulated and top 20 most significant down regulated proteins (p<0.05) shared between Batch 1 and Batch 2. F. IPA pathway analysis showing top 12 upregulated and top 12 down regulated pathways shared in both Batch 1 and 2 between *APOE4* and *APOE3* iHLCs. G. STRING network analysis of the most significant upregulated and down regulated proteins shared between Batch 1 and Batch 2 in *APOE4* versus *APOE3* iHLCs. H. Cellular component Gene Ontology (GO) analysis of the top 20 upregulated and top 20 down regulated proteins in *APOE4* versus *APOE3* iHLCs. n=5 per group. Data are shown as mean ±SD. ns=not significant, ***p<0.001.

### APOE4 isogenic iHLCs display impaired mitochondrial function and increased glycolysis

We next examined mitochondrial and glycolytic function in both isogenic pairs of iHLCs based on proteomic data. *APOE4* iHLCs derived from both Pair A and B isogenics had significantly reduced basal, maximal, proton leak, and ATP-production linked respiration (**Figure 5A-D**). *APOE4* iHLCs also showed reduced complex IV driven respiration/flux in both isogenic pairs, with disparate results for complex I (reduced in group B) and complex II (increased in group A) fluxes (**Figure 5E and F**). Mitochondrial membrane potential and mitochondrial superoxide production were increased in Pair A *APOE4* iHLCs but not Pair B cells (**Figure 5G-I, L-N**). Hydrogen peroxide (H_2_O_2_) levels, a form of ROS, were increased in both Pair A and B *APOE4* iHLCs (**Figure 5J and O**). Mitochondrial calcium (Ca^2+^) levels were increased in Pair B *APOE4* iHLCs but not Pair A (**Figure 5K and P**).

**Figure 5.**
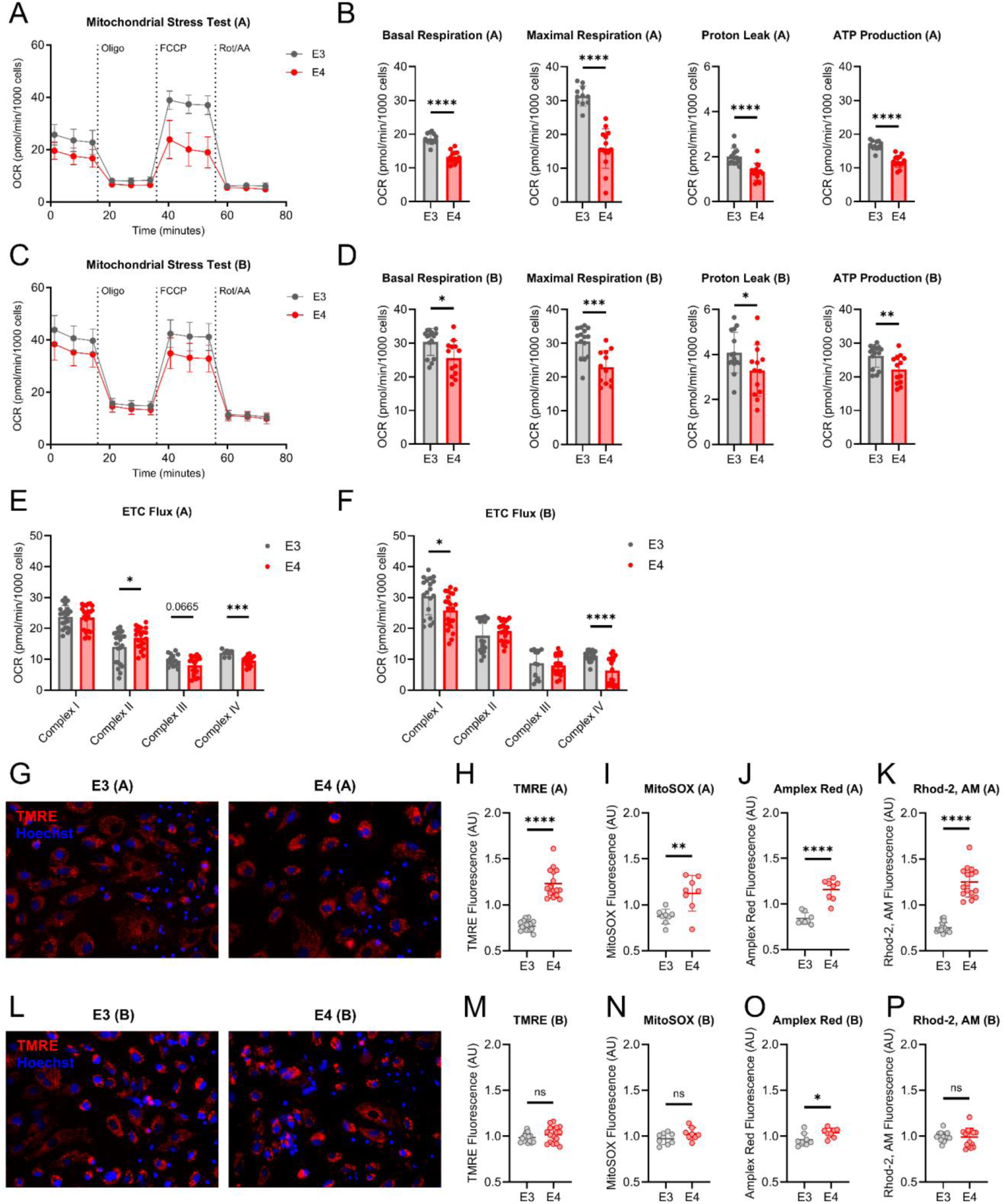
Mitochondrial function in *APOE* isogenic iHLCs. A. Mitochondrial stress test (MST) tracing from isogenic Pair A. B. Quantification of basal respiration, maximal respiration, proton leak, and ATP-production linked respiration from isogenic Pair A MST. C. MST tracing from isogenic Pair B. D. Quantification of basal respiration, maximal respiration, proton leak, and ATP-production linked respiration from isogenic Pair B MST. E. Electron transport chain (ETC) flux through complex I, II, III, and IV in isogenic Pair A. F. ETC flux through complex I, II, III, and IV in isogenic Pair B. G. Representative TMRE images from isogenic Pair A. H. TMRE, I. MitoSOX, J. Amplex Red, K. Rhod-2, AM fluorescence intensities from isogenic Pair A. L. Representative TMRE images from isogenic Pair B. M. TMRE, N. MitoSOX, O. Amplex Red, P. Rhod-2, AM fluorescence intensities from isogenic Pair B. Data are shown as mean ±SD. ns=not significant, *p<0.05, **p<0.01, ***p<0.001, ****p<0.0001.

Glycolytic function was increased in both isogenic pairs of iPSC-derived *APOE4* iHLCs, including basal glycolysis and glycolytic capacity (**Figure 6A-D**). Based on this data we wanted to examine glucose/pyruvate oxidation. Using UK5099, a mitochondrial pyruvate carrier (MPC) inhibitor, we found that mitochondrial pyruvate oxidation was reduced/impaired in both pairs of *APOE4* isogenic iHLCs as indicated by a smaller acute response (**Figure 6E-H**) (31). Unlike *APOE3* iHLCs, *APOE4* cells treated with UK5099 also exhibited lower maximal respiration compared to untreated *APOE4* cells, suggesting an increased reliance on glucose/pyruvate metabolism (**Figure 6E-H**). Unsurprisingly, glucose metabolism associated proteins were largely upregulated in proteomic analysis (**Figure 6I and J**).

**Figure 6.**
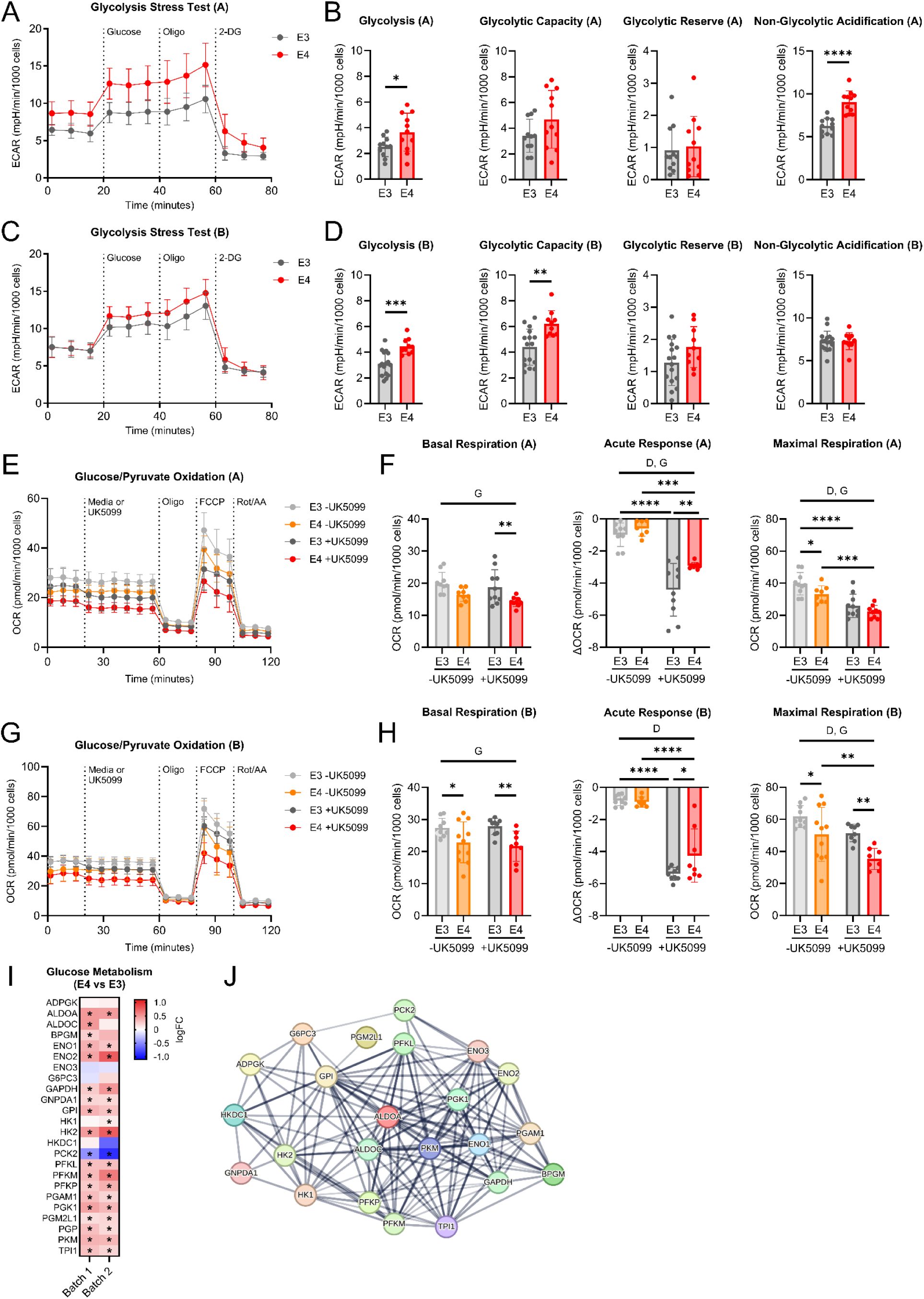
Glycolytic function and glucose/pyruvate oxidation in *APOE* isogenic iHLCs. A. Glycolytic stress test (GST) tracing from isogenic Pair A. B. Quantification of basal glycolysis, glycolytic capacity, glycolytic reserve, and non-glycolytic acidification from isogenic Pair A GST. C. GST tracing from isogenic Pair B. D. Quantification of basal glycolysis, glycolytic capacity, glycolytic reserve, and non-glycolytic acidification from isogenic Pair B GST. E. Glucose/pyruvate oxidation stress test from isogenic Pair A. F. Quantification of basal respiration, acute response, and maximal respiration from isogenic Pair A glucose/pyruvate oxidation stress test. G. Glucose/pyruvate oxidation stress test from isogenic Pair B. H. Quantification of basal respiration, acute response, and maximal respiration from isogenic Pair B glucose/pyruvate oxidation stress test. I. Significant DE proteins involved in glucose metabolism pathway from IPA. J. Glucose metabolism protein network STRING analysis. Data are shown as mean ±SD. *p<0.05, **p<0.01, ***p<0.001, ****p<0.0001. D=drug, G=genotype.

### Cholesterol and lipid metabolism are altered by APOE genotype in iHLCs

We next wanted to examine long chain fatty acid (LCFA) oxidation in these iHLCs. This was done using Etomoxir (Eto) which is an inhibitor of carnitine palmitoyltransferase-1 (CPT1), a key rate limiting step in FAO (32). Interestingly, *APOE4* cells treated with Eto showed a reduced acute response in both isogenic pairs indicating impaired LCFA oxidation (**Figure 7A-D**). Additionally, maximal respiration was lower in *APOE4* iHLCs when treated with Eto, an effect not seen in *APOE3* cells (**Figure 7A-D**). This points towards *APOE4* iHLCs having an increased reliance for LCFAs. Finally, we performed lipid droplet (LD) staining and found that *APOE4* iHLCs have an increased number of LDs per cell, but the size of LDs was not changed (**Figure 7E-H**). The SREBP2 protein network, or sterol regulatory element-binding protein 2, which is a transcription factor critical for cholesterol homeostasis and synthesis was upregulated in *APOE4* iHLCs when compared to their *APOE3* counterparts (**Figure 7I**).

**Figure 7.**
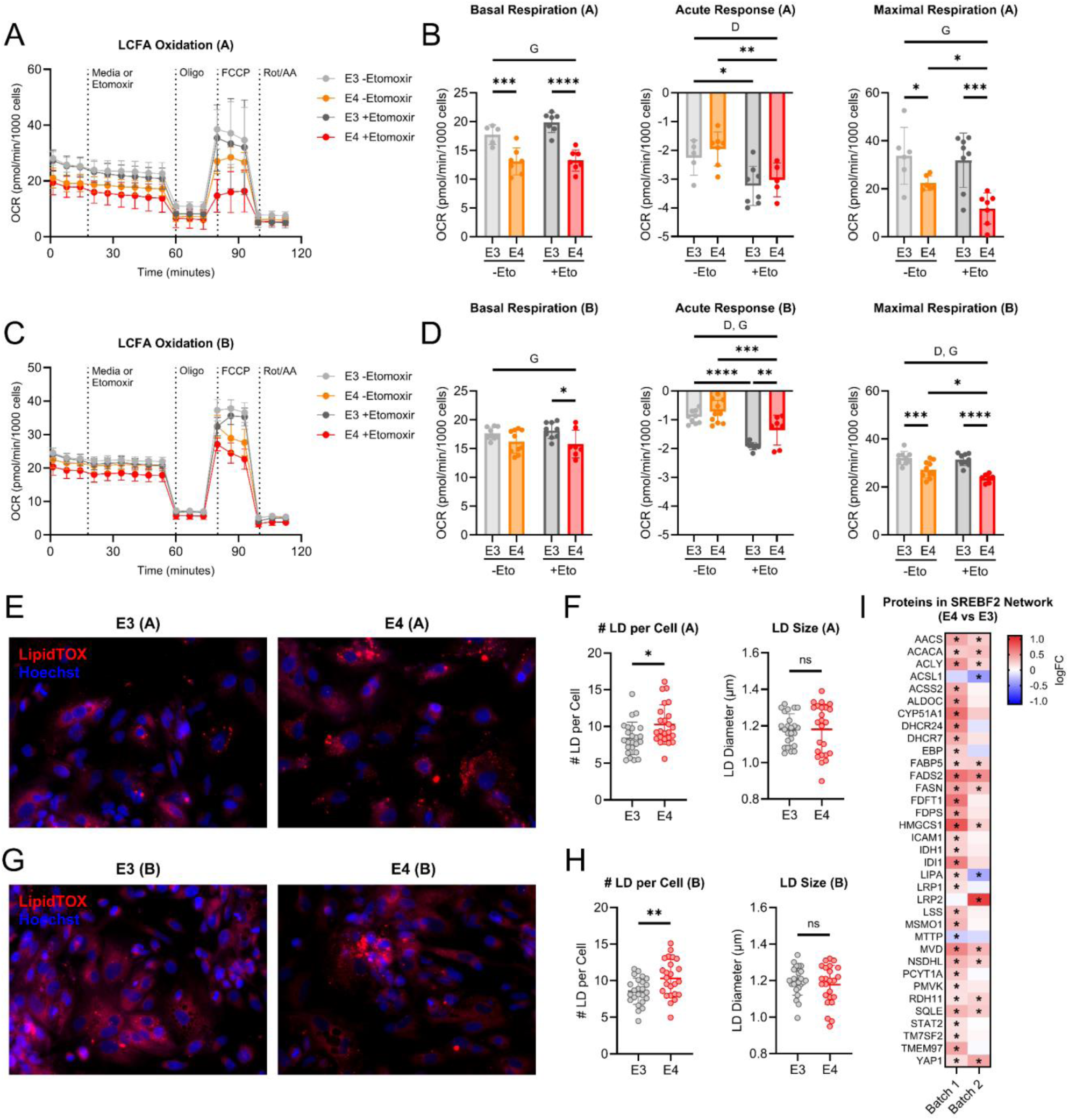
Fatty acid oxidation and LipidTOX staining in *APOE* isogenic iHLCs. A. Long chain fatty acid (LCFA) oxidation stress test from isogenic Pair A. B. Quantification of basal respiration, acute response, and maximal respiration from isogenic Pair A LCFA oxidation stress test. C. LCFA oxidation stress test from isogenic Pair B. D. Quantification of basal respiration, acute response, and maximal respiration from isogenic pair B LCFA oxidation stress test. E. Representative LipidTOX images from isogenic Pair A. F. Quantification of lipid droplet (LD) numbers per cell and diameter (μm) from isogenic Pair A. G. Representative LipidTOX images from isogenic Pair B. H. Quantification of LD numbers per cell and diameter (μm) from isogenic Pair B. I. Comparison analysis showing significantly altered proteins in SREBF2 network from proteomics. Data are shown as mean ±SD. ns=not significant, *p<0.05, **p<0.01, ***p<0.001, ****p<0.0001. D=drug, G=genotype.

## Discussion

While much is known regarding the brain and *APOE* genetic variation, the relationship with hepatic health remains relatively understudied. Recent studies have begun to address this knowledge gap and increase our understanding of the importance *APOE* genotype may play in maintaining liver metabolism (16–18). Leveraging multiple approaches, we show that *APOE4* drives alterations to hepatic metabolism. We observed sex and genotype dependent changes to metabolism in the liver of young *APOE* TR mice and shifts in metabolism in *APOE* isogenic iHLCs.

When measuring hepatic APOE levels, we did not observe significant differences across sex or genotype. This finding is interesting, as other studies have reported similar findings or have shown that *APOE4* reduces hepatic APOE expression (17, 33). However, these discrepancies may be attributed to differences in mouse age and/or strain. Furthermore, changes to liver health observed in this study may be driven by APOE function rather than differences in protein levels.

We explored how *APOE* genotype might alter the hepatic proteome and mitochondrial function. Remarkably, *APOE4* females had a large number of differentially expressed proteins when compared to their *APOE3* counterparts, an effect less pronounced in males. This is a critical consideration since women with at least one *APOE4* allele are at a higher risk of developing AD than men (34, 35). Analysis of altered pathways in the whole liver revealed a genotype by sex interaction. Specifically, *APOE4* females exhibited an upregulation of the electron transport chain (ETC) and oxidative phosphorylation when compared to *APOE3* females, whereas *APOE4* males showed a down regulation of these pathways. These changes were relatively consistent with functional respiratory data where *APOE4* females showed increased respiration for multiple respiratory states when compared to *APOE3* females.

Examining differences between sexes revealed that *APOE3* male mice had an upregulation of not only oxidative phosphorylation, but peroxisomal lipid metabolism as well when compared to females. An upregulation of lipid metabolism at peroxisomes may explain increased lipid driven respiration in *APOE3* male mice. This is supported by past research showing peroxisome and mitochondrial lipid metabolism are coupled (36–38). While we performed respiration on isolated mitochondria, this increase in peroxisomal lipid metabolism may have primed mitochondria in male mice. We also found that proteins associated with lipid metabolism were differentially expressed across all comparisons. This includes an increase of farnesyl-diphosphate farnesyltransferase 1 (Fdft1) in *APOE4* versus *APOE3* female mice. An increase in FDFT1 was also seen in *APOE4* iHLCs when compared to *APOE3* cells. This protein is involved in the mevalonate pathway and has been shown to be previously modified by *APOE* genotype in astrocytes and oligodendrocytes (39, 40). Finally, acetyl-CoA acetyltransferase 2 (Acat2), a protein involved in the biosynthesis of cholesterol, is down regulated in *APOE4* female and male mice. Past research has shown that ablation of *Acat2* can lead to increased plasma triglycerides (TGs) and cause mice on a high fat diet to become susceptible to insulin resistance, while *Acat2* overexpression promotes cholesterol metabolism and reduces blood glucose levels (41–43).

In addition to changes in mitochondrial energy and lipid metabolism, we also observed decreased glycolysis in *APOE4* females when compared to their *APOE3* counterparts. This a notable finding as the liver plays a major role in glucose metabolism (44). Interestingly, a down regulation of Rap1 signaling in *APOE4* versus *APOE3* male mice was observed. Numerous studies have shown that an upregulation of Rap1 signaling can protect the liver from hepatic steatosis and has also implicated it in glucose metabolism/signaling (45, 46). Sex-specific differences were also observed in the TCA cycle, with *APOE4* female mice exhibiting downregulation of this pathway, while *APOE4* male mice showed upregulation compared to their APOE3 counterparts. This is interesting because state 3S respiration is increased in *APOE4* female mice, an effect not seen in *APOE4* males. Sex comparisons revealed that *APOE3* male mice actually had a decrease in the TCA cycle when compared to their female counterparts. This may indicate increased enzymatic activity since *APOE3* males had higher pyruvate/malate driven respiration. Though, this requires further investigation into succinate dehydrogenase activity.

We also examined proteins involved in mitochondrial import between whole liver and isolated mitochondria in female mice. Specifically, there was an upregulation of the mitochondrial protein import pathway (Timm and Tomm machinery) in the whole liver of *APOE4* female mice. However, when we examine isolated mitochondria, we saw the opposite. This discrepancy could indicate several possibilities, including impaired import of mitochondrial proteins, selection of mitochondria less efficient at protein import during isolation, or potential compensation in *APOE4* female mice.

*APOE* isogenic iHLCs were used to examine hepatocyte-specific changes. Several upregulated proteins in *APOE4* iHLCs have been previously linked to AD and extracellular matrix (ECM) remodeling (19, 47–54). This includes proteins such as membrane metalloendopeptidase or neprilysin (MME), decorin (DCN), and F-spondin/spondin-1 (SPON1). Notable proteins decreased in *APOE4* iHLCs included APP, ferritin heavy chain 1 (FTH1), insulin-like growth factor binding protein 7 (IGFBP7), SPARC related modular calcium binding 2 (SMOC2), and metalloproteinase inhibitor 3 (TIMP3). One of these studies specifically investigated *APOE4* dependent changes in serum proteomics from AD patients and found an increase in SPON1 (19). However, this change was independent of the *APOE* genotype. Similar to our findings, they also observed alterations to ECM proteins. This is a notable finding as ECM proteins play a critical role in liver fibrosis (55, 56).

Proteins and pathways associated with metabolism were altered in *APOE4* iHLCs. This included a down regulation to pathways associated with mitochondrial function. This data is further supported by decreased mitochondrial respiration and ETC function in *APOE4* iHLCs. Unsurprisingly, *APOE4* iHLCs had increased H_2_O_2_ levels, and an upregulation of pathways known to be associated with mitochondrial stress such as cGAS-STING signaling (57). Interestingly, there was upregulation of glycolysis and associated proteins including hexokinase 2 (HK2) and SLC2A1 (glucose transporter type 1 or GLUT1) in *APOE4* cells. Glycolytic function was increased in APOE4 iHLCs, reflecting the elevated demand for glucose and pyruvate as fuel substrates. This is an interesting finding as there was a decrease in proteins and pathways associated with glycolysis/glucose metabolism in the livers of mice. This difference may be explained by the fact that iHLCs are isolated cells, in contrast to whole tissue, which contains numerous cell types with potentially different signaling dynamics. Irrespective of this, past studies have shown that *APOE* genotype can modulate glycolysis and glucose metabolism (58–60). Additionally, *APOE4* iHLCs have increased demand for fatty acids. Lipid droplet (LD) accumulation is also higher in *APOE4* cells indicating altered lipid homeostasis. Looking further at proteomics we found that proteins in the SREBP2 network are upregulated indicating increased cholesterol and fatty acid synthesis (61). Taken together these results indicate dysregulated lipid metabolism in *APOE4* iHLCs.

This study had several key strengths including the use of *APOE* isogenic iHLCs to resolve hepatocyte specific changes. We also showed sex and *APOE* genotype dependent changes in mice at a young age (4 months) which is comparable to a young adult (20-30 years of age in humans) (62). However, we did not use female iHLCs which would be valuable to examine sex differences in hepatocytes specifically. Finally, the impact of *APOE* genotype is not limited to *APOE3* and *APOE4* homozygosity, as these do not represent the full range of allelic combinations found in the population. Future studies may benefit from examining the *APOE2* allele in relation to hepatic health.

In conclusion, this study demonstrates that *APOE* genetic variation significantly impacts hepatic health, altering mitochondrial function as well as glucose and lipid metabolism. These findings raise additional questions including the role liver metabolism and health could contribute to AD risk and onset. This highlights the need for follow-up studies to validate and expand upon these results.

## Acknowledgements and Funding

This study was supported by the Margaret “Peg” McLaughlin and Lydia A. Walker Opportunity Fund, the University of Kansas Alzheimer’s Disease Center P30AG072973 (HMW, JKM, and JPT), Kansas Center for Metabolism and Obesity Research Center P20 GM144269 (JPT), R01AG078186 (HMW), R00AG056600 (HMW), R01DK121497 supplement (JPT), R01AG069781 (JPT), VA Merit Review grant 1I01BX002567 (JPT), Alzheimer Association Grant 23AARG-1023294 (HMW), T32HD057850 (VC), T32 Brain Health (CRL and CNJ). The content is solely the responsibility of the authors and does not necessarily represent the official views of the National Institutes of Health.

## Author Contributions

Acquisition of Funding (HMW, JKM), Project Conceptualization (CRL, HMW), Investigation (CRL, CNJ, VC, EF, MB, HMW), Resources (JPT, PCG, JKM, HMW), Methodology (CRL, JPT, HMW) Supervision (HMW), Writing – Original Draft Preparation (CRL, HMW), Writing – Review & Editing (All Authors)

---

**Supplemental Figure 1.**
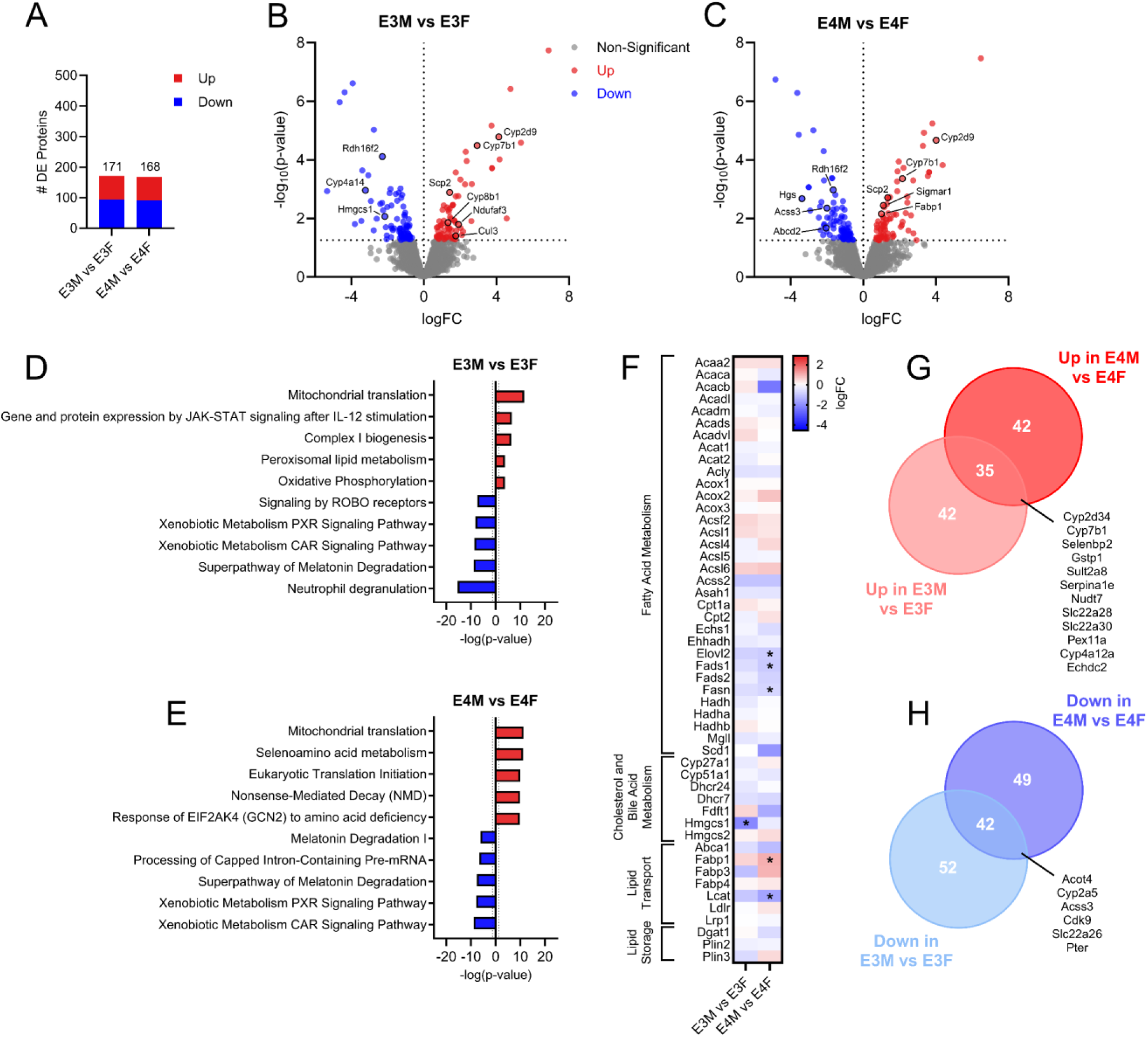
Whole liver proteomics showing sex-specific differences. A. DE proteins between male and female mice of the same genotype. B. Volcano plot of up and down regulated proteins between male and female *APOE3* mice. C. Volcano plot of up and down regulated proteins between male and female *APOE4* mice. D. Up and down regulated pathways between male and female *APOE3* mice. E. Up and down regulated pathways between male and female *APOE4* mice. F. Fatty acid metabolism, cholesterol/bile acid metabolism, lipid transport, and lipid storage pathway protein expression patterns between male and female mice. G. Upregulated proteins shared between male mice regardless of genotype. H. Down regulated proteins shared between male mice regardless of genotype. n=3 for proteomic outcomes.

**Supplemental Figure 2.**
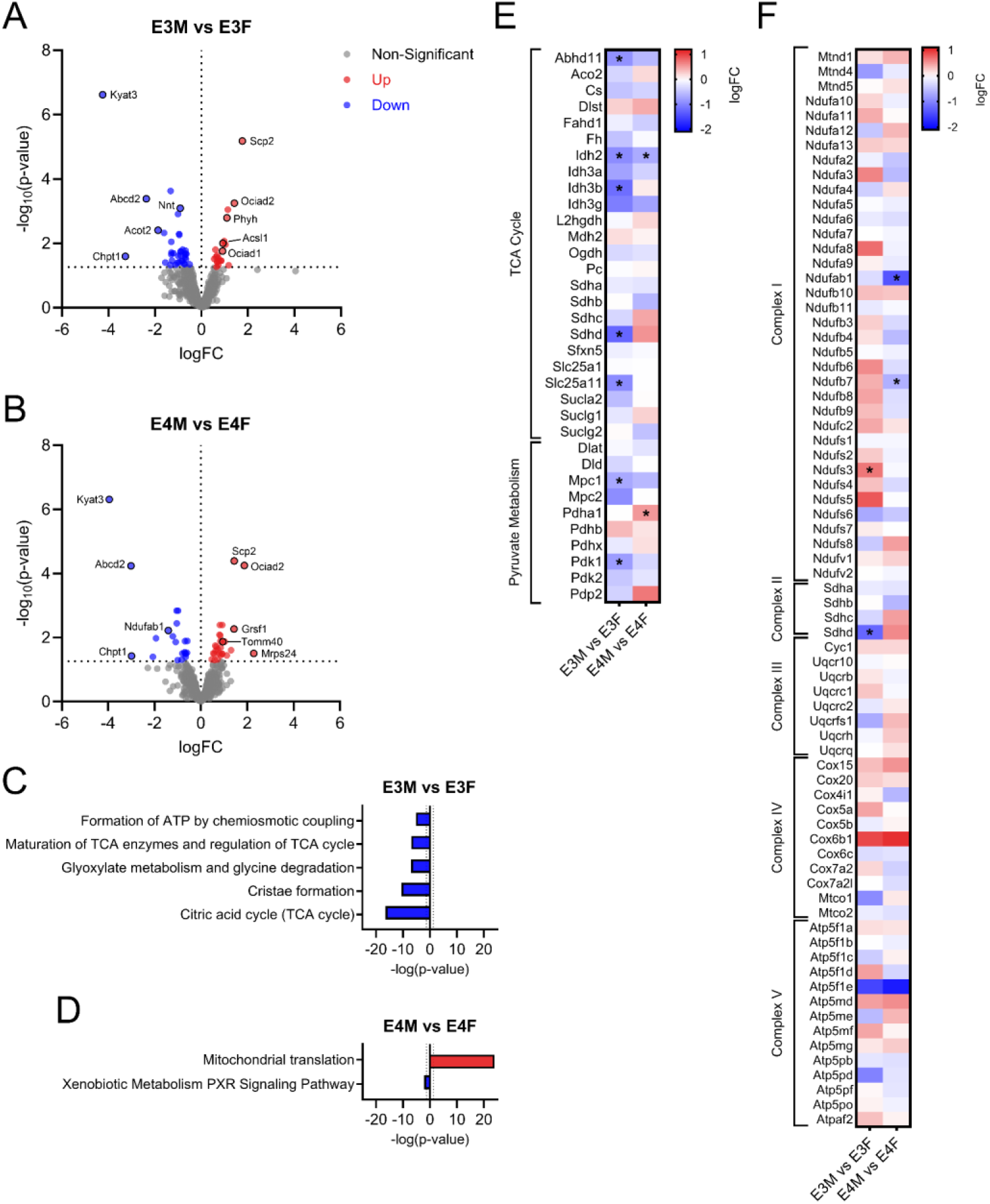
Isolated mitochondrial liver proteomics showing sex-specific differences. A. Volcano plot of up and down regulated proteins between male and female *APOE3* mice. B. Volcano plot of up and down regulated proteins between male and female *APOE4* mice. C. Up and down regulated pathways between male and female *APOE3* mice. D. Up and down regulated pathways between male and female *APOE4* mice. E. TCA cycle and pyruvate metabolism pathway protein expression patterns between male and female mice. F. Oxidative phosphorylation complex I, II, III, IV, and V component expression patterns between male and female mice. n=3 for proteomic outcomes.

**Supplemental Figure 3.**
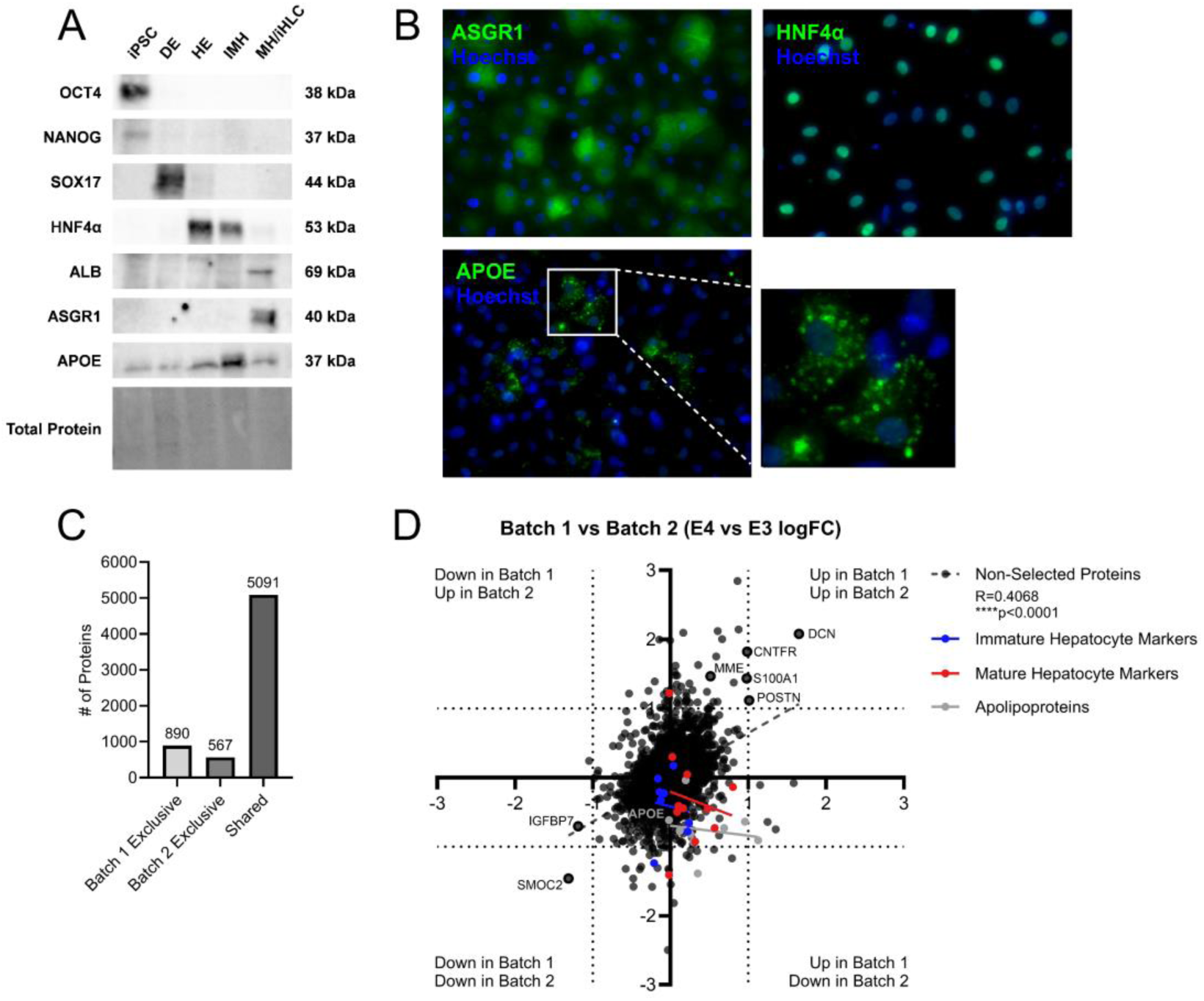
iHLC characterization and differentiation proteomics quality control. A. Western blot (WB) analysis of protein expression for iPSC (OCT4, NANOG), definitive endoderm (SOX17), early/immature hepatocyte (HNF4α), and mature hepatocyte (ALB, ASGR1, APOE) markers. B. Representative images showing expression of ASGR1 and APOE in iHLCs. C. Comparison of proteins identified in Batch 1, Batch 2, or both batches of proteomic samples. D. Pearson correlation analysis of protein expression patterns between Batch 1 and Batch 2 proteomic samples.

